# Genetic and cellular basis of impaired phagocytosis and photoreceptor degeneration in CLN3 disease

**DOI:** 10.1101/2024.06.09.597388

**Authors:** Jimin Han, Sueanne Chear, Jana Talbot, Vicki Swier, Clarissa Booth, Cheyenne Reuben-Thomas, Sonal Dalvi, Jill M. Weimer, Alex W. Hewitt, Anthony L. Cook, Ruchira Singh

## Abstract

**Purpose:** CLN3 Batten disease (also known as Juvenile Neuronal Ceroid Lipofuscinosis; JNCL) is a lysosomal storage disorder that typically initiates with retinal degeneration but is followed by seizure onset, motor decline and premature death. Patient-derived CLN3 disease iPSC-RPE cells show defective phagocytosis of photoreceptor outer segments (POSs). Because modifier genes are implicated in CLN3 disease, our goal here was to investigate a direct link between *CLN3* mutation and POS phagocytosis defect.

**Methods:** Isogenic control and *CLN3* mutant stem cell lines were generated by CRISPR-Cas9-mediated biallelic deletion of exons 7 and 8. A transgenic *CLN3*^Δ^*^7-8/^*^Δ^*^7-8^* (*CLN3*) Yucatan miniswine was also used to study the impact of *CLN3*^Δ^*^7-8/^*^Δ^*^7-8^* mutation on POS phagocytosis. POS phagocytosis by cultured RPE cells was analyzed by Western blotting and immunohistochemistry. Electroretinogram, optical coherence tomography and histological analysis of *CLN3*^Δ^*^7/8^* and wild-type miniswine eyes were carried out at 6-, 36-, or 48-month age.

**Results:** *CLN3*^Δ^*^7-8/^*^Δ^*^7-8^* RPE (*CLN3* RPE) displayed reduced POS binding and consequently decreased uptake of POS compared to isogenic control RPE cells. Furthermore, wild-type miniswine RPE cells phagocytosed *CLN3*^Δ^*^7-8/^*^Δ^*^7-8^* POS less efficiently than wild-type POS. Consistent with decreased POS phagocytosis, lipofuscin/autofluorescence was decreased in *CLN3* miniswine RPE at 36 months-of-age and was followed by almost complete loss of photoreceptors at 48 months of age.

**Conclusions:** *CLN3*^Δ^*^7-8/^*^Δ^*^7-8^* mutation (that affects up to 85% patients) affects both RPE and POSs and leads to photoreceptor cell loss in CLN3 disease. Furthermore, both primary RPE dysfunction and mutant POS independently contribute to impaired POS phagocytosis in CLN3 disease.

## Introduction

CLN3 disease is the most common form of Batten disease (neuronal ceroid lipofuscinoses or NCLs) that develops in early childhood. Symptoms include progressive loss of vision and neurodegeneration of the brain, ultimately leading to premature death^1,2^. Most affected CLN3 disease patients (∼75%) have a 966 bp deletion that entirely removes exon 7 and 8 from the *CLN3* transcript^3^. Despite this, there is significant variability in the disease phenotype of CLN3 disease patients, including those harboring the same common *CLN3* mutation^4–6^. The lack of genotype-phenotype relationship in CLN3 disease has suggested a contribution of modifier genes to the disease development and progression^4^.

Murine models have been widely used as an *in vivo* model of CLN3 disease^7–11^. However, despite some evidence of molecular and pathological alterations, there is poor recapitulation of retinal degeneration phenotype in these models^12–14^. To overcome the lack of visual degeneration of the existing CLN3 models, a transgene miniswine model was developed that carries the orthologous common 966 bp deletion found in most human patients^96^. Importantly, *CLN3^Δ7–8/Δ7-8^* Yucatan miniswine (*CLN3* miniswine) shows both progressive neuronal loss, motor dysfunction, and vision impairment phenotypes similar to those seen in individuals witch CLN3 disease^15^.

Although retina degeneration is a prominent pathology in CLN3 disease, there is limited knowledge of the disease mechanism(s) underlying retinal degeneration in CLN3 disease. Clinical and histopathologic studies on CLN3 disease retina have shown autofluorescent lipopigment buildup in neurons with degeneration of several retinal cell type(s), including the retinal pigment epithelium (RPE)^16–18^. Furthermore, histopathologic analyses of CLN3 disease donor eyes showed decreased autofluorescent material, lipofuscin, in the RPE and in contrast elevated autofluorescence in the photoreceptor cell layer^16,19–21^. We have recently replicated decreased RPE autofluorescence in the CLN3 disease donor eyes in patient-derived human induced pluripotent stem cell (hiPSC)- RPE cells (harboring the common 966 bp deletion) post- chronic daily exposure to a physiological load of photoreceptor outer segments (POSs)^20^. However, because this study compared CLN3 disease patient-derived hiPSC-RPE cells to non- isogenic control hiPSC-RPE cells, it is unclear if the observed POS phagocytosis defect in CLN3 disease is a direct consequence of *CLN3* mutation. Note that, as previously mentioned modifier genes have strongly implicated in molecular alterations observed in CLN3 disease^4^.

CRISPR-based gene editing has been routinely used for *in vitro* correction of genetic mutations in human pluripotent stem cells, including hiPSCs and human embryonic stem cells (hESCs)^22–24^. In fact, CRISPR-based gene editing of hiPSCs and hESCs have been routinely utilized to decipher genotype-phenotype relationship in several retinal diseases^25–28^. For example, we have previously used gene-corrected hiPSC lines from patients with Doyne honeycomb retinal dystrophy (DHRD) to show that R345W mutation in EFEMP1 is causal for drusen accumulation in DHRD^29^. Similarly, CRISPR-Cas9 mediated repair *of CLCN2* mutation rescued dysfunctional POS phagocytosis by iPSC-RPE cells from patients with *CLCN2*-associated retina degeneration^30^. Also, Neural retina-specific leucine zipper protein (NRL)-deficient hESC-retina organoids engineered via CRISPR-Cas9 gene editing showed a direct role of NRL in rod photoreceptor specification^31^. Overall, gene-editing strategies including both correction of genetic defect in patient hiPSCs and introduction of disease-associated mutation in control hESC/hiPSCs provides a suitable approach to definitively link the independent contribution of genetic variants to molecular and pathological defects in a human-based cellular model.

Therefore, to link disease-causing *CLN3* mutation to the observed impaired phagocytosis in CLN3 disease, here, we used CRISPR/Cas9 gene editing to generate isogenic H9 hESC lines carrying homozygous deletion of exons 7 and 8 in CLN3 (referred to here in as *CLN3^Δ7-8/Δ7-8^*). Additionally, we utilized a large animal model of CLN3 disease to study the impact of common 966 bp *CLN3* deletion mutation on POS phagocytosis and autofluorescence/lipofuscin accumulation.

## Materials and Methods

### Availability of data and materials

Request for original data files and reagents and materials used in this study will be fulfilled by contacting the corresponding author, Dr. Ruchira Singh.

### Procurement and use of hESC line

H9 hESCs were obtained from WiCell and used under approval from the Tasmania Health and Medical Human Research Ethics Committee (#13502) and Institutional Regulatory Board (IRB) (RSRB00056538)at the University of Rochester and conformed with the ethical norms and the declaration of Helsinki.

### Animals

WT and transgenic (*CLN3^Δ7-8/Δ7-8^*) miniswine^8^ used in these studies were obtained from Exemplar Genetics and all animal studies were conducted following the approval of the Institutional Animal Care and Use Committees at Exemplar Genetics (Protocol # MRP2018-004).

### CRISPR-Cas9-mediated editing of *CLN3*

Editing of *CLN3* to delete exons 7 and 8 was done as previously described^32,33^. Briefly, 8 x 10^5^ H9 hESCs in single-cell suspension were resuspended in 100 µL electroporation buffer from the Human Stem Cell Nucleofector Solution 2 kit (Lonza, VPH-5022). Alt-R Cas9 Electroporation enhancer (1.1 µM) (IDT, #1075915) and assembled CRISPR/Cas ribonucleoprotein consisting of 1.4 µM dual crRNA (*Table 1*) (IDT) and 1.2 µM Alt-R S.p. Hifi Cas9 Nuclease V3 (IDT, #1081060) were mixed with cells and transferred to a nucleofection cuvette. Electroporation was conducted using Amaxa Nucleofector 2b Device (Lonza) with Nucleofector program B-016. Electroporated cells were seeded at low density (2,000 cells/well) in a Matrigel (Corning, FAL354277)-coated 6 well-plates with CloneR (STEMCELL Technologies, #05889)/ mTeSR^TM^1 media (STEMCELL Technologies, #85850). When single-cell colonies were approximately 200 µm in diameter, each colony was picked and seeded into 2 wells of duplicate Matrigel-coated 96 well-plates with one plate for cryopreservation and the other for genotyping.

### PCR screening and genotyping

When cells from the genotyping 96-well plate were 50-60% confluent, DNA extraction was done using QuickExtract DNA Extraction Solution (Epicentre, #E09050) according to manufacturer’s instructions. Subsequently, 50 ng of DNA extract was amplified through PCR reaction using KAPA HotStart PCR Kit, with dNTPs (Millenium Science, ROC-07958897001), and CLN3 F1 and CLN3 R1 primers that span the targeted region (*Table 1*).

### Characterization of pluripotency

Embryonic stem cell colonies at 80% confluency were dissociated with ReLeSR (STEMCELL Technologies, #05872). Cell aggregates were transferred into ultra-low attachment 6-well plates in Complete KSR EB medium (KnockOut™ SR (Life Technologies, #10828-028) 20%, DMEM/F-12, GlutaMAX™ supplement (Life Technologies, #10565018) 79%, MEM Non-Essential Amino Acids solution 1% (Life Technologies, #11140050), 2-mercaptoethanol 0.1 mM (Sigma, M6250) for embryoid body (EB) formation. Medium was refreshed every 2 days. On day 4, EBs were plated onto Matrigel-coated wells for further differentiation in Complete KSR EB medium for 16 days. Pluripotency markers were analyzed by quantitative real-time PCR (qRT-PCR) and immunocytochemistry.

For qRT-PCR analyses, total RNA was isolated from positive clones with the RNeasy Plus Mini Kit (Qiagen, #74134) according to the manufacturer’s protocol. Reverse transcription of 1 μg RNA was performed using Omniscript RT kit (Qiagen, #205111). CLN3 transcript was amplified, and PCR products were sequenced with primers, CLN3 F2 and CLN3 R2 that span exons 6-9 (***Table 1***).

For immunocytochemical analyses, hESC colonies were passaged and cultured on Matrigel-coated coverslips in mTeSR1 medium until 60% confluency. Cells were then fixed with 4% paraformaldehyde in PBS for 20 minutes at room temperature. Fixed cells were incubated for one hour in blocking solution (5% fetal bovine serum (FBS) in PBS and 5% FBS with 0.1% Triton X-100 in PBS for extracellular and intracellular markers respectively) followed by incubation with the primary antibodies for NANOG, OCT4, TRA-1-60 and SSEA4 overnight at 4°C. This was followed by incubation in host-specific secondary antibody for one hour at room temperature and thereafter mounting and visualization by fluorescent microscopy. Further details of primary antibody concentration and source can be found in ***Table 2***.

### Virtual karyotyping

Genomic DNA (200 ng) from isogenic H9 *CLN3^Δ7-8/Δ7-8^*cell lines were analyzed for copy number variation using Illumina HumanCytoSNP-12 beadchip array. B allele frequency (BAF) and log R ratio (LRR) of each single nucleotide polymorphism (SNP) marker were collected from GenomeStudio (Illumina, USA). These were used for CNV analyses which was performed using PennCNV^34^ and QuantiSNP^35^ with default parameter settings. Genomic regions having at least 20 contiguous single nucleotide polymorphisms (SNPs) or genomic regions with SNPs spanning at least 1 MB were designated as chromosomal copy number variation^36^.

### RPE differentiation and culture from hESCs

Differentiation of hESC to RPE was performed as previously described^20,37^. Briefly, embryoid bodies (EBs) were generated from colonies of hESCs cultured in either mTeSR™ or mTeSR™ (Stemcell technologies). On day 6 of EB culture, EBs were plated onto laminin-coated tissue culture plates and fed with neural induction medium or NIM whose composition is described in^20,37^. The cell culture medium was switched to retinal differentiation medium (RDM) on day 14. RDM composition is described in^20,37^. hiPSC-RPE cells were dissected in patches from adherent cultures around day 60-90 when characteristic RPE morphology could be observed. The RPE patches at passage P0 were seeded and passaged onto either again a non-permeable plastic 24-well plate or onto transwell (Corning) insert (0.4 µm pore size). RPE cells at < P4 were used in all experiments.

### Primary miniswine RPE culture

Primary miniswine RPE cultures were performed as previously described^39,40^. Briefly, after enucleation, eyes were placed in 0.2 % povidone-iodine solution for 10 minutes on ice followed by 5 rinses in 1000U/mL penicillin-streptomycin solution. The anterior portion of the eye and the retina was next removed (note the retina was used for POS isolation) and eyecups were transferred to 6-well plates filled with 1ml of prewarmed 0.5% trypsin with 5.3 mM EDTA in HBSS without calcium and magnesium. Subsequently, the 6 well plates were placed in a 37°C incubator for 30 minutes. At the end of the incubation, RPE cells were collected in a 15mL tube containing prewarmed media (1X DMEM with 4.5 g/L glucose, L-glutamine, and sodium pyruvate and 10% FBS) and gently centrifuged ( 300 rcf) to collect the RPE cell pellet. The RPE cell pellet was resuspended and plated in a T25 flask in culture media (1X DMEM with 4.5 g/L glucose, L-glutamine, and sodium pyruvate, 1% NEAA, 1% penicillin/streptomycin) containing 10 %FBS. Once the RPE cells have reached confluence, the FBS concentration in the culture media was reduced to 1% and subsequently RPE cells were passaged and replated onto laminin-coated 24-wells or transwells.

### POS phagocytosis assay

POS phagocytosis assay was performed as previously described20. Bovine POSs were obtained commercially from InVision BioResources (Cat. #98740, Seattle, WA) and porcine POSs prepared from wild-type and CLN3 miniswine as previously described^38–40^. Note that POS was isolated eyes from three WT and four CLN3 miniswine eyes of ∼30-36 months of age.

For evaluating POS uptake, as described earlier^20^, mature monolayer of RPE cell in culture were fed unlabeled POS or FITC-labeled POS (∼10-40 POS/RPE cell)^2^ for two hour at 37°C with 10% FBS media supplementation. Thereafter, to remove POS (unbound) on the RPE cell surface, RPE cells were thoroughly washed with 1X PBS. Subsequently, RPE cells were either harvested for Western blotting or processed for immunocytochemistry experiments. Quantitative analysis of the number of FITC-POS phagocytosed included a consideration of FITC-particle size. Specifically, a threshold of <5µm allowed us to evaluate the amount bound, ingested POS but eliminate aggregated POS from the analyses.

For evaluating POS binding and POS internalization, we used a previously published protocol^41^ that was also utilized to evaluate POS binding versus internalization in CLN3 disease hiPSC- RPE^20^. Briefly, the position of FITC-particles relative to ZO1 was used to evaluate the amount of bound versus internalized FITC-POS particles^42^. Specifically, to analyze the relative position of the FITC-POS within the cell, FITC-POS fed RPE cells post-immunocytochemical labeling with ZO1 antibody were imaged as confocal z-stacks spanning the entire height of the RPE cells. The images were subsequently analyzed with ImageJ software to determine the number of apical versus basal FITC-POS particles relative to ZO1.

### RPE Autofluorescence measurements

As previously described^20,43^, for *in vitro* experiments evaluating autofluorescence accumulation, RPE cells were fed POS daily (∼20-40/RPE cell/day) for 14 days. At day 14, to remove any unbound POS, RPE monolayer was thoroughly washed with 1X PBS. RPE cells were then immediately fixed and processed by immunocytochemistry where the red channel (ex: 546 nm and em: 560-615 nm) was exclusively used for evaluation of RPE autofluorescence without any antibody staining. Quantitative analyses of autofluorescent particles were performed per DAPI-stained cells for count and per viewing area for area covered by the autofluorescent particles using ImageJ/FIJI and Microsoft Excel software.

For analysis of autofluorescence in miniswine eye samples, miniswine RPE whole mounts were utilized. Miniswine eyes were injected with 4% PFA following enucleation for 1h prior to dissection to remove the anterior portion and vitreous. The remaining retina/RPE eyecup was submerged in 4% PFA for a fixation of at least 4h at 4°C. Following two 1X PBS washes, samples were prepared for wholemount staining by carefully removing the RPE/choriocapillaris from the sclera. The RPE wholemount was then permeabilized/blocked in blocking buffer (1% bovine serum albumin (ImmunoReagents, Inc.), 0.2% Triton-X-100, 0.2% Tween-20) for 1h room temperature. Subsequently RPE Samples were washed and incubated with DAPI dye (Hoechst 33342; Life Technologies) for 15-30min at room temperature. After one last wash with 1X PBS, RPE samples were mounted onto slides with Prolong Gold (Life Technologies P36930) and cover slipped and imaged by confocal microscopy in the red channel (ex: 546 nm and em: 560-615 nm).

### Western blot

Western blotting was performed as described^20^. Briefly, RPE cells were lysed in Radio- Immunoprecipitation Assay (RIPA) buffer containing protease inhibitor cocktail (Sigma-Aldrich). Subsequently, protein quantification was performed with Pierce BCA assay and equal amount of protein was loaded for each sample and resolved on 4-20% Tris-HCl gradient gels, followed by transfer onto Polyvinylidene Fluoride (PVDF) membranes. The PVDF membrane was subsequently blocked with 5% dry milk in 1X PBS or commercially bought blocking buffer (LI-COR) for 1h at room temperature followed by primary antibody incubation overnight at 4°C. After primary antibody incubation, PVDF membrane was washed with 0.1% Tween-PBS and incubated in host-specific HRP or IRDye-conjugated secondary antibody (1:10,000) in 5% dry milk 0.1% Tween-PBS for 1h at room temperature. Azure C500 (Azure Biosystem) and Image Studio were used for analysis and quantification of the Western blot images. Note that all primary antibodies used for Western blotting analyses are listed in *Table 2*.

### Immunocytochemistry

Immunocytochemical analysis was performed as previously described^20,43^. Briefly, RPE on transwells were permeabilized and blocked in blocking buffer (10% normal donkey serum, 0.1% Triton-X-100) for 1h at room temperature. This was followed by overnight incubation in primary antibody in 0.5X blocker buffer at 4°C. Next day, RPE in transwells were washed and subsequently incubated with host-specific Alexa-conjugated secondary antibody (1:500) in 0.5X blocking buffer for 1h at room temperature. The RPE in transwells were next incubated with Hoechst 33342 (Life Technologies) for 15 min to stain the nuclei, then washed in 1X PBS for 5 min. Subsequently, transwell membranes were cut out and mounted onto slides with Prolong Gold (Life Technologies) and imaged with a confocal microscope (LSM 510 META, Zeiss, or Eclipse Ti2, Nikon). Confocal microscopy images were analyzed with ImageJ/FIJI software. All primary antibodies used can be found in *Table 2*.

### Histological analyses of miniswine retina

Whole eyes were fixed for ∼3 weeks in 10% neutral buffered formalin. Subsequently, the retina was dissected and the midperiphery region of the retina was used for processing and analyses.

Specifically, the dissected retina tissue was further fixed at room temperature in 4% PFA for 1h. After 1X PBS washes, the retina was processed for plastic embedding (Technovit 7100 kit; Electron Microscopy Sciences) as previously described^15^. Retina sections were cut at 3 μm thick and dried on a slide warmer. Multiple Stain Solution (Polysciences, Inc.) was used to stain the retina sections on slides, which were then mounted with permount (Fisher Scientific) and imaged by light microscopy.

### Transmission electron microscopy analyses

Transmission electron microscopy (TEM) was performed as previously described^20^. Briefly, RPE grown on transwells were fixed overnight in a solution containing 0.1 M sodium cacodylate, 2.5% glutaraldehyde and 4% paraformaldehyde. Next, RPE samples on transwell membrane were embedded in epoxy resin and processed to obtain 60 nm thick RPE section, then imaged at the Electron Microscopy Core at the University of Rochester using a Hitachi H-7650 Transmission Electron Microscope.

### TER measurements

EVOM2 volt-ohm meter (World Precision Instruments) was used to measure TER of RPE grown in transwell inserts as described^44^ using an). A TER measurements were reported as resistance per area or Ω.cm^-2^ after blank subtraction,

### ERG and OCT measurements

ERG was performed using RET*eval*^®^ device (LKC technologies) under anesthesia as previously described ^15^. Briefly for scotopic ERG measurements, animals were dark adapter for 20 mins.

Photopic ERG measurements were conducted under standard illumination with exposure to room light for over 10 minutes. Scotopic and photopic a-wave and b-wave amplitudes data from each eye for each analysis was measured separately but graphical representation of an individual measurement (e.g., scotopic a wave) shows averaged data for both eyes together.

Retinal layer thickness of miniswine via OCT was assessed using the Leica instrument spectral-domain OCT system (Bioptigen Envisu R2200; Leica, Danaher, Washington, DC). Prior to imaging, eyes were dilated to increase the field of view using topical tropicamide 1% and ophthalmic 2.5% phenylephrine eye drops. For each eye, high density scans consisted of 1000x100x1 (A scans x B scans x repeated B scans) for averaging, with a 12mm x 12mm volume scan centered upon the retina superior to the optic nerve. Multiple images were obtained from each eye for analysis. OCT analysis to access retinal layer thickness was performed on 3 scans per eye using Bioptigen InVivoVue and ImageJ software

Experimental set-up, data analyses and statistical testing.

For all experiments comparing isogenic control and *CLN3* RPE (with the exception of immunocytochemistry for RPE signature proteins), parallel age-matched cultures of RPE monolayer in transwell inserts that displayed TER > 150 Ω.cm^-2^ were utilized. Data is presented as mean ± sem. Two-tailed unpaired student’s t-test was used to compute significance and Microsoft Excel was used to plot the graphs. For quantitative analysis of confocal microscopy data, at least 5 distinct images (different viewing areas) per sample were used in each experiment. In each individual figure n refers to the biological replicate sample size for data from either isogenic control versus *CLN3^Δ7-8/Δ7-8^* hESC-RPE cells derived from distinct EB differentiations, or *ex vivo* or *in vivo* analyses of age-and sex matched wild-type and *CLN3* miniswine retina and RPE.

## Results

### Generation and characterization of isogenic control and CLN3^Δ*7-8/*Δ*7-8*^ (*CLN3*) hRPE cells

To generate *CLN3^Δ7-8/Δ7-8^* isogenic cell lines, we utilized a well-characterized hESC line, H9^45^. *CLN3^Δ7-8/Δ7-8^* intron 6 and the other within intron 8 which are spaced 968 bp apart in the *CLN3* gene (***Figure 1a***). Our first attempt of CRISPR editing yielded a H9 *CLN3^+/^*^Δ*7-8*^ cell line with monoallelic deletion of exons 7-8 in CLN3 gene (***Figure. 1b***). To obtain a biallelic deletion of exons 7-8 in the *CLN3* gene, we repeated CRISPR editing on the H9 *CLN3^+/^*^Δ*7-8*^ cell line. Subsequent PCR revealed two truncated DNA bands (***Figure. 1b***), which despite having different deletion sizes had complete deletion of exons 7-8 from both CLN3 alleles. The successful biallelic deletion of exons 7-8 was confirmed through sequencing of the *CLN3* transcript using cDNA and primers targeting regions outside exons 7-8 in the homozygous H9 *CLN3^Δ7-8/Δ7-8^* cell line (***Figure. 1c***).

**Figure 1.**
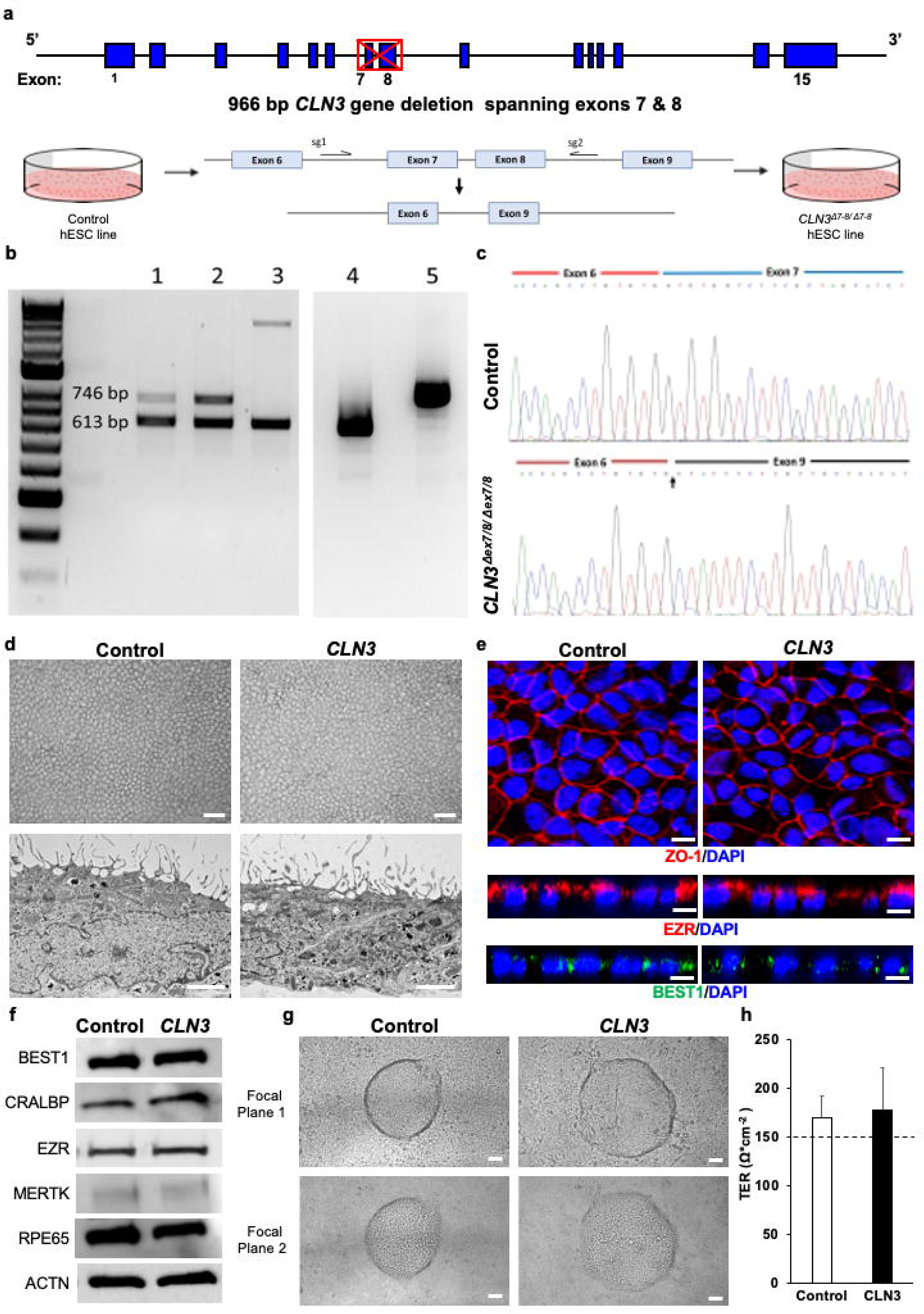
Generation and characterization of isogenic control and CLN3 hESC RPE. **a)** Schematic depicting CRISPR-Cas9 deletion of exons 7 and 8 of CLN3 in H9 hESC. Note the CLN3 locus and the location of dual guides to excise exons 7-8 from the CLN3 gene. **b)** Identification of exons 7-8 deletion in CLN3 gene in positive clones. Genomic PCR of clones with homozygous deletion (lane 1 and 2) and heterozygous deletion (lane 3). RT-PCR analysis of homozygous 1kb deletion clone (lane 4) and control (lane 5). **c)** Sequencing alignment of cDNA for control (top panel) and *CLN3^Δ7-8/Δ7-8^* (bottom panel) cell lines shows successful biallelic deletion of exons 7-8 reflected as splicing of exon 6 to exon 9 (indicated by an arrow). **d)** Representative light microscopy images (top panel) and electron microscopy images (bottom panel) showing the expected RPE morphology and apical microvilli in isogenic control and *CLN3^Δ7-8/Δ7-8^* hESC RPE (*CLN3* hRPE) (scale bar = 100µm). **e)** Representative confocal microscopy images showing similar and expected localization of tight junction protein, ZO1 (red, top panel), RPE microvilli protein, EZR (red, middle panel), and basolaterally expressed RPE protein, BEST1 (green, bottom panel) in control and *CLN3* hRPE cells. Nuclei were stained with DAPI (blue) (scale bar = 10µm). **f)** Representative Western blot images showing expected presence of RPE signature proteins in both control and CLN3 hRPE cells. ACTN served as the loading control in these experiments. **g)** Representative light microscopy images showing evidence of transepithelial fluid flux with presence of fluid domes in polarized monolayers of control and *CLN3* hRPE cells. Note the presence of RPE both within the fluid dome (focal plane 1, top panel) and outside the fluid dome (focal plane 2, bottom panel) (scale bar = 50 µm). **h)** Transepithelial resistance (TER) measurements showing evidence of a tight epithelial monolayer in both control and *CLN3* hRPE cultures. Note the dotted line represent the known TER threshold for RPE cells in *vivo*^48^. n ≥ 3 for all experiments in figure 1

Genome-wide copy number variation analysis demonstrated that the H9 *CLN3^Δ7-8/Δ7-8^* isogenic cells have normal diploid karyotypes (46,XX). No chromosomal alterations were detected in the edited cells when compared to the unedited control cells based on analysis of the B allele frequency and Log R ratio (***Figure. S1***).

Pluripotency of isogenic H9 and H9 *CLN3^Δ7-8/Δ7-8^* cell lines was confirmed by positive reactivity for pluripotency markers OCT4, NANOG, TRA-1-60, and SSEA4 (***Figure. S2***). Quantitative real-time polymerase chase reaction (qRT-PCR) analysis of embryoid bodies also demonstrated expression of genes specific for all three germ layer markers (endoderm, ectoderm, and mesoderm) (***Figure. S2***).

Isogenic control and *CLN3^Δ7-8/Δ7-8^* H9 ESC lines successfully differentiated to RPE cells using our previously described protocols^20,29,37,46^. Light microscopy and transmission electron microscopy (TEM) of control RPE and *CLN3^Δ7-8/Δ7-8^* RPE (referred henceforth to as *CLN3* hRPE) showed the expected cell morphology including presence of elongated RPE microvilli in both control and *CLN3* hRPE cultures (***Figure 1d***). Confocal microscopy imaging post- immunocytochemical analyses also showed similar and expected localization of tight junction protein, ZO1, in both control and *CLN3* hRPE cultures (***Figure 1e***). Similarly, consistent with polarized expression of specific proteins in RPE cells, orthogonal view of confocal microscopy images of control and *CLN3* hRPE showed apical localization of microvilli protein, EZR, and basolateral localization of BEST1 (***Figure 1e***). Qualitative Western blotting analyses showed robust presence of RPE signature proteins, BEST1, CRALBP, EZR, MERTK, RPE65, as well as loading control (ACTN) in control and *CLN3* hRPE cells. In addition, in agreement with transepithelial fluid movement and formation of a polarized epithelial monolayer^20,47^, light microscopy images showed presence of fluid domes in both control and *CLN3* hRPE cultures (***Figure 1f***). Lastly, transepithelial epithelial resistance (TER) recordings showed formation of a tight epithelial barrier with both control and *CLN3* hRPE developing physiological TER of ∼150 Ω*cm^-2^ expected for RPE monolayer *in vivo*^48^.

Overall, CRISPR-Cas9 editing, we were able to generate pluripotent isogenic control and *CLN3^Δ7-8/Δ7-8^* H9 hESC lines. Notably, isogenic control versus *CLN3* hRPE cells showed similar morphology with presence of several RPE signature proteins and the expected RPE cell polarity and tight junction characteristics akin to native human RPE *in vivo*.

Disease causing *CLN3* mutation is independently sufficient to promote impair POS binding and consequently impair POS phagocytosis by RPE cells.

We have previously shown reduced POS binding and consequently impaired POS phagocytosis by patient-derived CLN3 disease hiPSC-RPE cells compared to control hiPSC-RPE cells^1^. To link CLN3 function to POS phagocytosis by RPE cells, we evaluated uptake of non-diseased POS by isogenic control versus *CLN3* hRPE cells.

Confocal microscopy imaging post-immunocytochemical analyses of control and *CLN3* mutant RPE cells that were fed FITC-labeled POS (∼20-40 POS/RPE cell) for 2h showed fewer number of phagocytosed POSs in *CLN3* hRPE cells compared to parallel cultures of control RPE cells (***Figure 2a-2c***). Autofluorescence accumulation (lipofuscin) in RPE cells is a consequence of POS phagocytosis and we have previously been able to show disease-associated reduction in lipofuscin (autofluorescence) in CLN3 disease patient-derived hiPSC-RPE cells after chronic (14 day) daily supplementation of a physiological dose of POS by measuring autofluorescence levels in the red channel (ex: 546 nm and em: 560-615 nm). Consistently, confocal microscopy analyses of autofluorescence levels in the red channel in POS-fed RPE cells (∼40 POS per day for 14 days) showed decreased accumulation of autofluorescent material (count, area) in the *CLN3* hRPE cells compared to control RPE cells (***Figure 2d-2f***).

**Figure 2.**
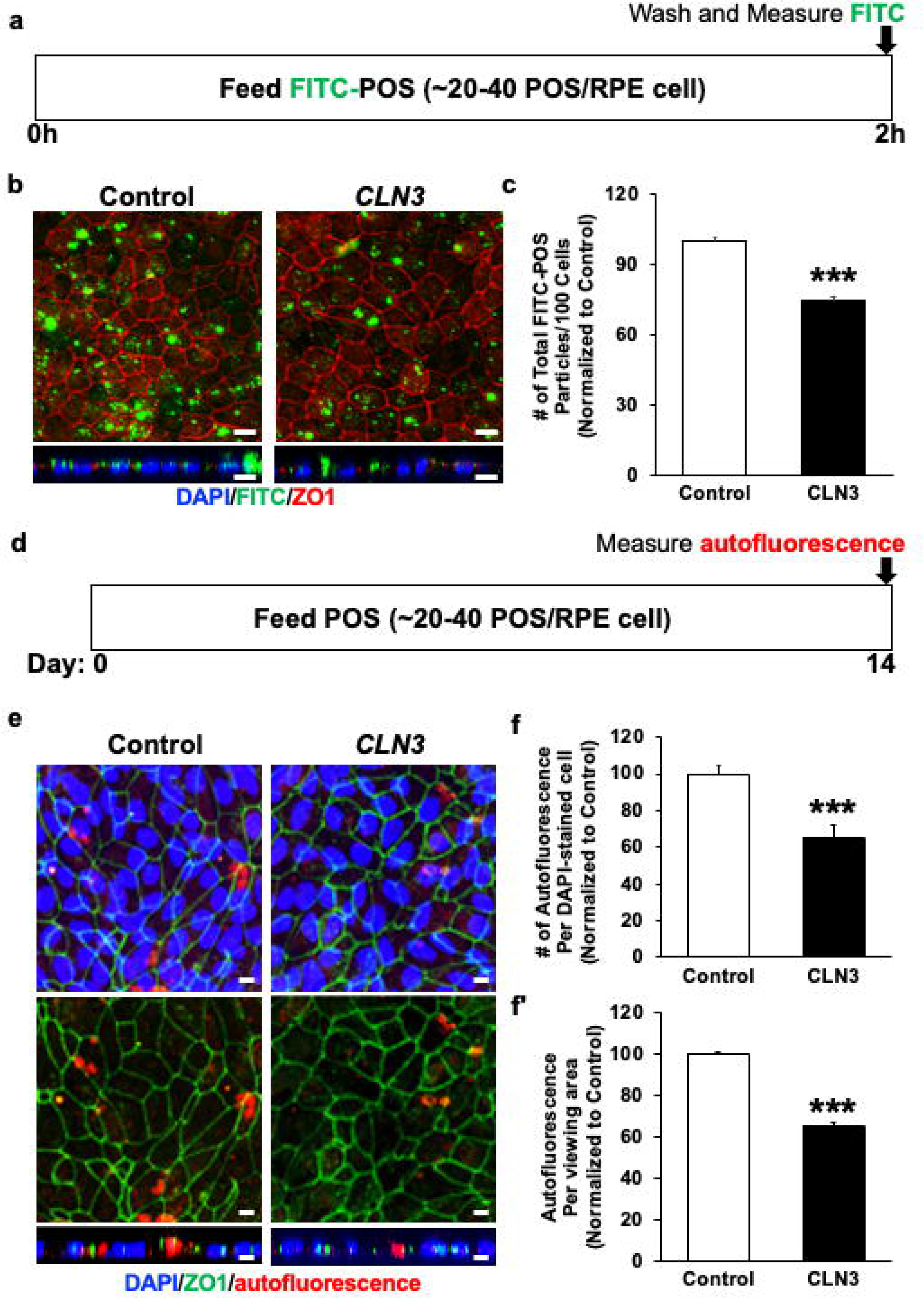
Evaluation of POS phagocytosis and RPE autofluorescence by control versu *CLN3* hRPE cells. **a)** Schematic of experimental assay utilized to measure POS uptake by control and *CLN3* hRPE cells after feeding a physiologic dose of POS (∼20-40 POS/RPE cell) for 2h. **b, c)** Representative confocal microscopy images (b, scale bar 10 µm) and quantitative analyses (c) showing decreased phagocytosis of FITC-POS particles (green, panel b) but similar localization of tight junction protein (ZO1) in *CLN3* hRPE cells compared to control hRPE cells. Nuclei were stained with DAPI (blue) (scale bar = 10µm). ***p ≤ 0.001. **d)** Schematic of experimental assay utilized to measure RPE autofluorescence after daily chronic POS feeding (∼20-40 POS/RPE cell) for a duration of 14 days by control and *CLN3* hRPE cells after feeding a physiologic dose of POS (∼20-40 POS/RPE cell) for 2h. **e, f)** Representative confocal microscopy images (e, scale bar 10µm) and quantitative analyses (f, f’) showing decreased RPE autofluorescence (count, area) in the red channel (ex: 546 nm and em: 560-615 nm) but similar localization of tight junction protein (ZO1) in *CLN3* hRPE cells fed daily with POS compared to control hRPE cells fed daily with POS. Nuclei were stained with DAPI (blue) (scale bar = 10µm). ***p ≤ 0.001. n ≥ 3 for all experiments in figure 2

To confirm that reduced POS phagocytosis by *CLN3* mutant RPE cells is a direct consequence of defective POS binding by *CLN3* mutant RPE cells, we next utilized a previously published protocol^20,41^ to analyze bound versus internalized FITC-labeled POS (∼20-40 POS/RPE cell) post-2h POS feeding in isogenic control versus *CLN3* hRPE cells (***Figure 3***). Compared to parallel cultures of control RPE cells, *CLN3* mutant hRPE that displayed decrease in total POS uptake (***Figure 2b, 2c and 3a***) also showed decrease in both the amount of apically localized (bound) and basally localized (internalized) FITC-POS particles relative to ZO1 (***Figure 3b-3e***). However, no difference in the internalized FITC-POS relative to bound FITC-POS was seen in control versus *CLN3* hRPE (***Figure 3f***).

**Figure 3.**
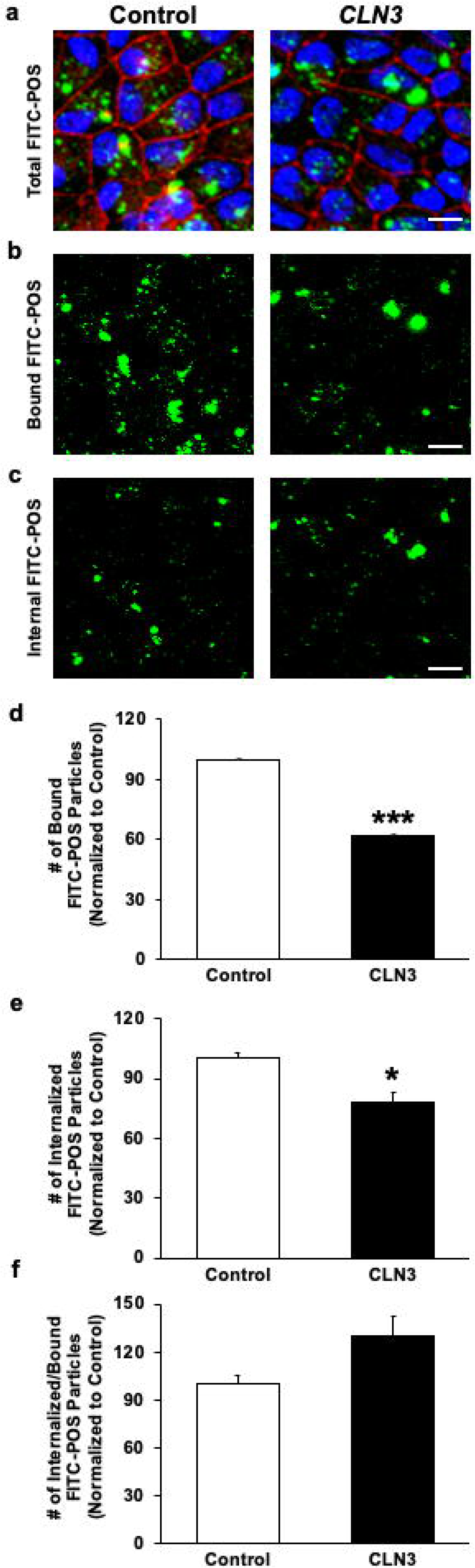
Evaluation of POS binding versus internalization by control versus *CLN3* hRPE cells. **a-c)** Representative confocal images of FITC-POS fed (2h) RPE monolayers showing decreased number of total FITC-POS particles (a), apical FITC-POS compared to ZO1 (consistent with reduced FITC-POS binding (b), and basal FITC-POS compared to ZO1 (consistent with reduced FITC-POS internalization) (c) in *CLN3* hRPE cells compared to control hRPE cells. Note that a published protocol that utilized position of ZO-1 relative to FITC-POS to estimate bound versus internalized POS was used in these experiments (scale bar = 10 μm). d-f) Quantitative analyses showing decreased number of both bound FITC-POS (d) and internalized FITC-POS (e) in *CLN3* hRPE cells compared to control hRPE cells. Consistent with a POS binding defect, the number of internalized FITC-POS relative to number of bound FITC-POS was unchanged between control and *CLN3* hRPE cells (**f**). *p ≤ 0.05, ***p ≤ 0.001. n ≥ 3 for all experiments in figure 3

Overall, these data confirm that *CLN3* mutation is independently sufficient to promote impaired POS binding and consequently defective POS phagocytosis by CLN3 disease RPE cells.

### *CLN3* mutant POS are phagocytosed less efficiently by wild-type RPE cells

Our published paper^20^ and results presented in this study (***Figure 2, 3***) show that disease-causing CLN3 mutation leads to impaired POS phagocytosis by RPE cells. However, these data lack the consideration of diseased POS to the phagocytosis process. Note that POS disorganization is an early pathological hallmark of CLN3 disease retina in living human eye^2,16,49,50^. Furthermore, several studies have implicated inflammation and oxidative stress in CLN3 disease^51–56^ and it has previously been shown that oxidized POS and POS exposed to free radicals are phagocytosed less efficiently by RPE cells^57,58^. Therefore, we next investigated the phagocytosis of control versus *CLN3* mutant POS.

We chose to utilize *CLN3* mutant POS isolated from *CLN3^Δ7-8/Δ7-8^* Yucatan miniswine in these experiments as it is well-established that POS in stem cell-derived retina organoids lack reproducibility and consistency potentially due to the lack of RPE^59,60^. Parallel cultures of wild-type miniswine RPE primary were fed either unlabeled wild-type miniswine POS or *CLN3* miniswine POS (∼10 POS/RPE cell) for 2 h. Subsequently, phagocytosis of POSs was measured by quantitative Western blotting analyses measuring the levels of RHO, a POS-specific protein, within the wild-type miniswine RPE cells (***Figure 4a, 4b***).

**Figure 4.**
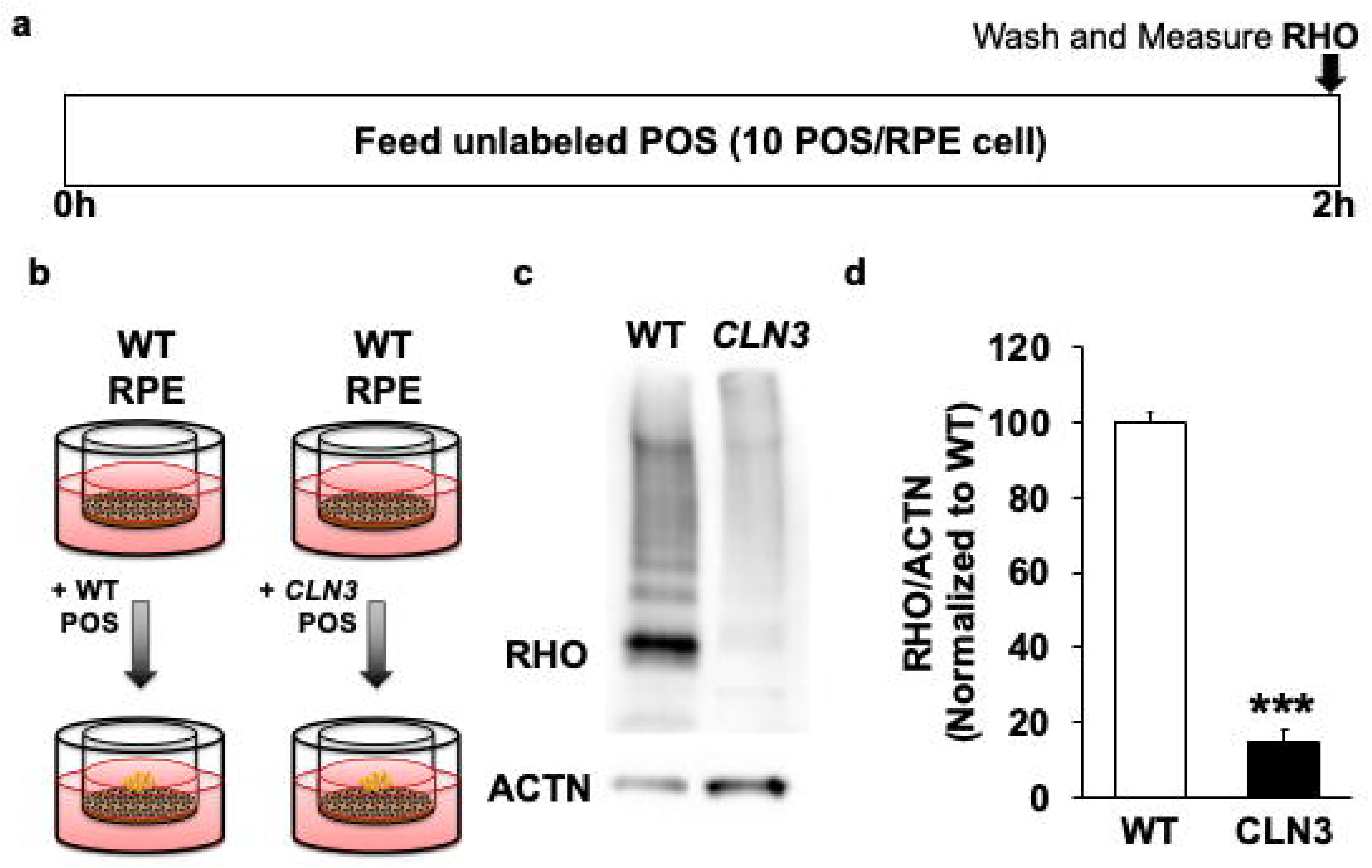
Evaluation of POS uptake of wild-type (WT) versus *CLN3* POS by RPE cells. **a, b)** Schematic of experimental assay utilized to measure POS phagocytosis by WT primary miniswine RPE post feeding of WT miniswine POS versus *CLN3* pig POS for 2h. **c, d)** Representative Western blot image (c) and quantification (d) showing decreased phagocytosis of *CLN3* miniswine POS by WT miniswine RPE compared to WT miniswine POS by WT miniswine RPE. ***p ≤ 0.001. n ≥3 for figure 4 experiment.

*CLN3* miniswine POS were phagocytosed less efficiently by wild-type miniswine RPE cultures compared to wild-type miniswine POS (***Figure 4c, 4d***). In fact, wild-type miniswine POSs were phagocytosed over 5-fold more efficiently than the *CLN3* miniswine POS (***Figure 4c, 4d***). These data suggests that CLN3 POS contribute to defective POS phagocytosis by RPE cells in CLN3 disease.

Disease-associated reduction in autofluorescence/lipofuscin levels precedes loss of outer nuclear layer in *CLN3* miniswine model.

Lipofuscin, the autofluorescence accumulation that accumulates in RPE cells because of incomplete digestion of POS by RPE cells, is reduced in CLN3 disease donor RPE cells^16,21^. CLN3 disease and CLN3 hRPE cells phagocytose (uptake) less POS and consequently have reduced RPE autofluorescence/lipofuscin compared to control RPE cells^20^ (***Figure 2e, 2f***). Thus, we utilized autofluorescence levels in the spectral wavelength consistent with lipofuscin, as a surrogate for efficacy of POS phagocytosis by RPE cells in *CLN3* miniswine retina.

Longitudinal analyses of autofluorescence levels (EX: EM) in the RPE cells of age- and sex-matched wild-type versus *CLN3* miniswine retina showed similar RPEautofluorescence in wild-type and *CLN3* miniswine retina at 6 months of age (***Figure 5a-5c***). In contrast, there was reduced lipofuscin in the *CLN3* miniswine RPE compared to wild-type miniswine RPE at 36 months of age (***Figure 5a-5c***). Note that in our previous study^15^, *CLN3* miniswine had shown reduced photopic and scotopic a wave at ∼30-36 months of age. ERG analyses of *CLN3* miniswine retina at 3 and 6 months of age, when there is no reduction in RPE lipofuscin/autofluorescence (***Figure 5a-5c***), showed similar photopic and scotopic a-wave amplitudes in wild-type and *CLN3* miniswine retina (***Figure S3***). Similarly, photopic and scotopic b-wave recordings were similar in wild-type and *CLN3* miniswine retina at 3 and 6 months of age (***Figure S3***).

**Figure 5.**
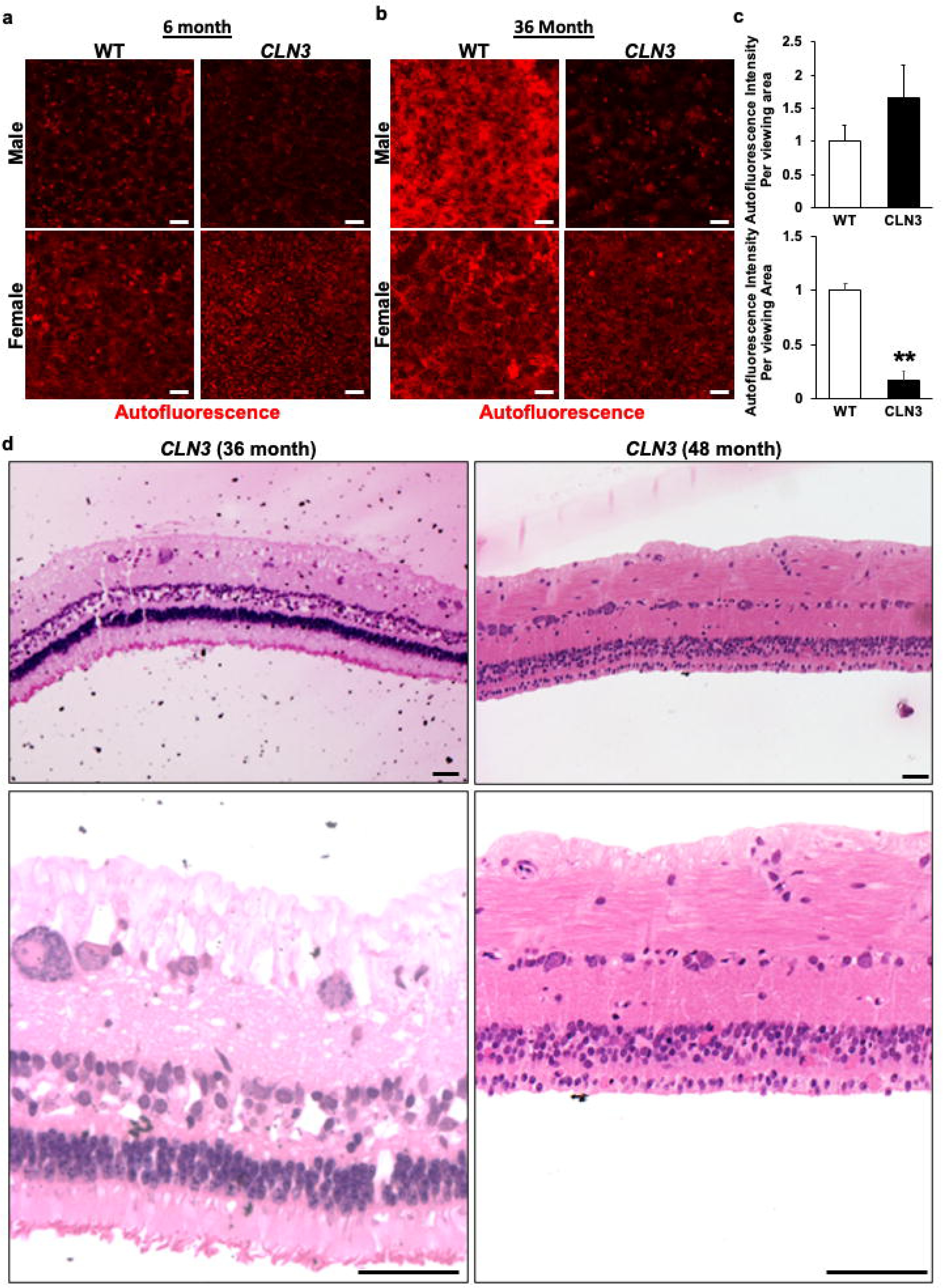
Longitudinal comparison of RPE autofluorescence levels and retina histology in *CLN3* miniswine eye. **a-c)** Representative confocal images (a, b) and quantitative analyses (c, c’) showing similar levels of autofluorescence accumulation in RPE of WT and *CLN3* miniswine at 6-month of age. In contrast, decreased autofluorescence accumulation was observed in *CLN3* miniswine RPE compared to WT miniswine RPE at the 36-months of age (scale bar 10 µm). **p ≤ 0.01. d) Representative images of *CLN3* miniswine retina sections at 36 and 48-months age showing presence of POSs at 36 months of age but absence of POSs with significant loss of the photoreceptor cell nuclei/ outer nuclear layer (ONL) at 48-months of age. n>3 for all experiments in figure 5. Scale bar = 5 μm.

Longitudinal histological analyses of *CLN3* miniswine retina at the 36- and 48-month timepoint showed the presence of POSs and multiple layers of outer nuclear layer (ONL) nuclei in the 36 month *CLN3* miniswine retina. In contrast, POSs were almost entirely gone with a near-complete loss of ONL in *CLN3* miniswine retina at 48 months of age. Note that optical coherence tomography (OCT) analyses of wild-type versus *CLN3* miniswine retina showed no significant changes in any of the retina cell layer including the ONL at 6 months of age, although the nerve fiber layer (NFL) was slightly reduced when measurements were taken 1 mm from the optic nerve (***Figure S4***).

Overall, these data show that reduction in RPE lipofuscin/autofluorescence coincides with photopic and scotopic visual deficits and precedes overt lack of POSs and ONL loss in *CLN3* miniswine retina.

## Discussion

CLN3 disease is the most common form of neuronal ceroid lipofuscinoses (NCLs). Apart from neurological symptoms, vision loss due to retinal degeneration is an early clinical hallmark of CLN3 disease. In a previous study, we used a CLN3 disease patient-derived hiPSC-RPE to show impaired POS phagocytosis^20^. Because gene modifiers have been strongly implicated in CLN3 disease^4^, we generated isogenic control and *CLN3* H9 hESC lines (hESCs; H9), as well as using *CLN3* miniswine model, to show that impaired POS phagocytosis is a direct consequence of disease-causing *CLN3* mutation (*CLN3^Δ7-8/Δ7-8^*), which contributes to reduced RPE lipofuscin/autofluorescence in CLN3 disease^20,49^.

Histopathological and clinical characterizations of CLN3 disease have suggested that retina degeneration starts at the POS^2,61^. Furthermore, CLN3 disease donor eyes show decreased RPE lipofuscin/autofluorescence^49^. Despite this, there have been limited investigation of POS phagocytosis in CLN3 disease^12,13,20^. This is presumably due to the postulation that photoreceptor cell death and the consequent lack of POS lead to reduced lipofuscin accumulation in RPE cells^49^. Here, we provide evidence that POS phagocytosis dysfunction and subsequently reduced RPE lipofuscin in CLN3 disease is a consequence of both primary RPE dysfunction (***Figure 2, 3***) and POS alterations (***Figure 4***). Specifically, by comparing the uptake of non-diseased POS by parallel cultures of isogenic control and *CLN3* hRPE cells (***Figure 2, 3***), we show that cell autonomous RPE dysfunction due to *CLN3* mutation (*CLN3*^Δ*7*^-8/^Δ*7-8*^*^-8^*) is sufficient for reduced POS phagocytosis in CLN3 disease. Similarly, by comparing the uptake of wild-type versus *CLN3* POS by wild-type RPE cells (***Figure 4)***, we show that POS alterations in CLN3 disease also independently contribute to decreased POS phagocytosis by RPE cells in CLN3 disease.

A limitation of our study is that we did not validate the role of POS alterations in impaired POS phagocytosis by RPE cells in the human-relevant hESC model. Similarly, we did not validate the POS phagocytosis defect *in vivo* in the *CLN3* miniswine model. This was due to consideration of the most optimal experimental approach and reduction in the use of tissue from a large animal model of the disease. The hESC-RPE, which have been consistently shown to possess properties of human RPE cells *in vivo*^62,63^, provided a suitable model system to investigate the ability of control versus *CLN3* RPE cells to phagocytose shed POSs. Similarly, the availability of *in vitro* protocols to assess POS binding versus internalization^20,41^ allowed us to interrogate the precise role of POS binding versus POS internalization in impaired POS phagocytosis by *CLN3* hRPE cells (***Figure 3***).

In contrast to RPE cells, the reproducibility of POS differentiation and maturation in the stem cell-derived retina organoid is variable^59,60^. Therefore, to avoid confounding variability due to experimental model system, we relied on primary POSs isolated from wild-type versus *CLN3* miniswine in experiments evaluating the role of diseased POS on the efficiency of POS phagocytosis by RPE cells in CLN3 disease. Note that recent advances in protocol for POS isolation from native retina allowed us to obtain sufficient POS from 3-4 miniswines. Furthermore, we were able to use RPE from the animals sacrificed for POS isolation to compare autofluorescence accumulation between wild-type versus *CLN3* miniswine RPE cells.

Reduced lipofuscin/autofluorescence accumulation preceded the loss of POSs and photoreceptors in the *CLN3* miniswine model and coincided with the earliest timepoint of scotopic and photopic visual deficit in *CLN3* miniswine (***Figure 5, S3***)^15^. This is consistent with pathological studies of human eyes that have shown neuronal depletion in CLN3 disease retina starting at the outer segments of the photoreceptors and proceeding inwards to the ganglion cell layer^2,61^. Apart from highlighting a central role of photoreceptor-RPE interaction in CLN3 disease (that needs further investigation), these data further highlight the value of *CLN3* miniswine model for studying the retina pathobiology of CLN3 disease.

## Conclusions

Using both *in vivo* (*CLN3* miniswine) and *in vitro* (isogenic hESC control and *CLN3* lines), we show that the most common disease-causing mutation in CLN3 disease (*CLN3*^Δ*ex7/*Δ*8/*Δ*ex7/*Δ*8*^) leads to POS alterations and primary RPE dysfunction that contribute to defective POS phagocytosis and reduced RPE autofluorescence/lipofuscin in CLN3 disease. Furthermore, reduced RPE autofluorescence/lipofuscin coincides with the earliest timepoint of scotopic and photopic visual deficit and precedes the loss of POSs and photoreceptors in the in a large animal model of CLN3 disease.

## Acknowledgments

We would like to acknowledge support of Exemplar Genetics staff in conducting studies (ERG, OCT, tissue recovery) that was performed at the Exemplar Genetics facilities.

## Funding

This work was supported by R01EY028167 (R.S.), R01EY030183 (R.S.), R01EY033192 (R.S.), funding from ForeBatten Foundation to both Singh and Weimer laboratory, Research to Prevent Blindness Unrestricted Challenge Grant to Department of Ophthalmology at University of Rochester and University Research Award given to R.S. by the University of Rochester. We would also like to acknowledge the Batten Disease Support and Research Association Australia (RGP204), assistance and Royal Hobart Hospital Research Foundation (17-205) to A.L.C. and A.W.H.

## Author’s contribution

Conceptualization and methodology: A.L.C., A.W.H., J.M.W.; R.S.

Experiment and analyses: A.L.C., A.W.H., C.B., C.R.T., J.H., J.M.W.; J.T., S.C., S.D., R.S.

Writing: Initial draft: R.S.

Writing – review & editing: A.L.C., A.W.H., C.B., C.R.T., J.H., J.M.W.; J.T., S.C., S.D., R.S.

Resources, A.L.C., A.W.H., J.M.W.; R.S.

## Notes

### Competing Interest Statement

The authors have declared no competing interest.

### Summary of Updates

The abstract (conclusions) has been updated

