## supplementary figure legends for "Genetic and cellular basis of impaired phagocytosis and photoreceptor degeneration in CLN3 disease"

**Figure S1. Copy number variation analysis of H9 *CLN3^Δ7-8/Δ7-8^* isogenic cell lines.**

**a. b)** Top and bottom panels show B allele frequency (BAF) and Log R ratio (LRR) respectively for the control (a) and ***CLN3^Δ7-8/Δ7-8^*** (b) hESC.

**Figure S2. Pluripotency characterization of isogenic *CLN3^Δ7-8/Δ7-8^* H9 human embryonic stem cell lines.**

**a)** Fluorescence microscopy images showing presence of pluripotency markers OCT4, NANOG, TRA-1-60 and SSEA4 in control and ***CLN3^Δ7-8/Δ7-8^***  hESCs (Scale bar =100 µm).

**b)** Graphical representation of gene expression levels of the pluripotency markers *OCT4* and *NANOG* and the germ layer markers PAX6 (ectoderm), GATA2 (mesoderm) and AFP (endoderm) in isogenic ***CLN3^Δ7-8/Δ7-8^*** embryoid bodies (EBs) relative to hESCs that were normalized to *EEF2* gene according to the ΔΔCt method.

**Figure S3. Scotopic and photopic ERG measurements in wild-type (WT) versus *CLN3* miniswine at 3 and 6 months of age.**

**a, b)** Graphical representation of ERG measurements showing similar amplitudes of photopic a-wave (a) and scotopic a-wave (b) in WT and *CLN3* miniswine retina at 3 (3M) and 6 months (6M) of age. Each individual dot represents a separate measurement.

**c, d)** Graphical representation of ERG measurements showing similar amplitudes of photopic b-wave (a) and scotopic b-wave (b) in WT and *CLN3* miniswine retina at 3 (3M) and 6 months (6M) of age. Each individual dot represents a separate measurement.

**Figure S4. OCT analyses of retina thickness in WT versus *CLN3* miniswine at 6 month of age.**

**a-c)** Quantitative analyses of retina thickness at a distance of 1mm (a), 3 mm (b) and 5 mm (c) from the optic nerve showed similar retinal thickness of individual retina cell layers between WT and *CLN3* miniswine at 6 months of age with the exception of nerve fiber layer (NFL) that was slightly thinner in *CLN3* miniswine retina at 1mm from the optic nerve. Individual parameters measured including thickness of total retina, outer nuclear layer (ONL), NFL, inner plexiform layer (IPL), outer plexiform layer (OPL) and outer retina layer (ORL). *p ≤ 0.05
