## Supplementary figures and images for "Genetic and cellular basis of impaired phagocytosis and photoreceptor degeneration in CLN3 disease"

### supplementary figures

#### Supplementary Figure 1

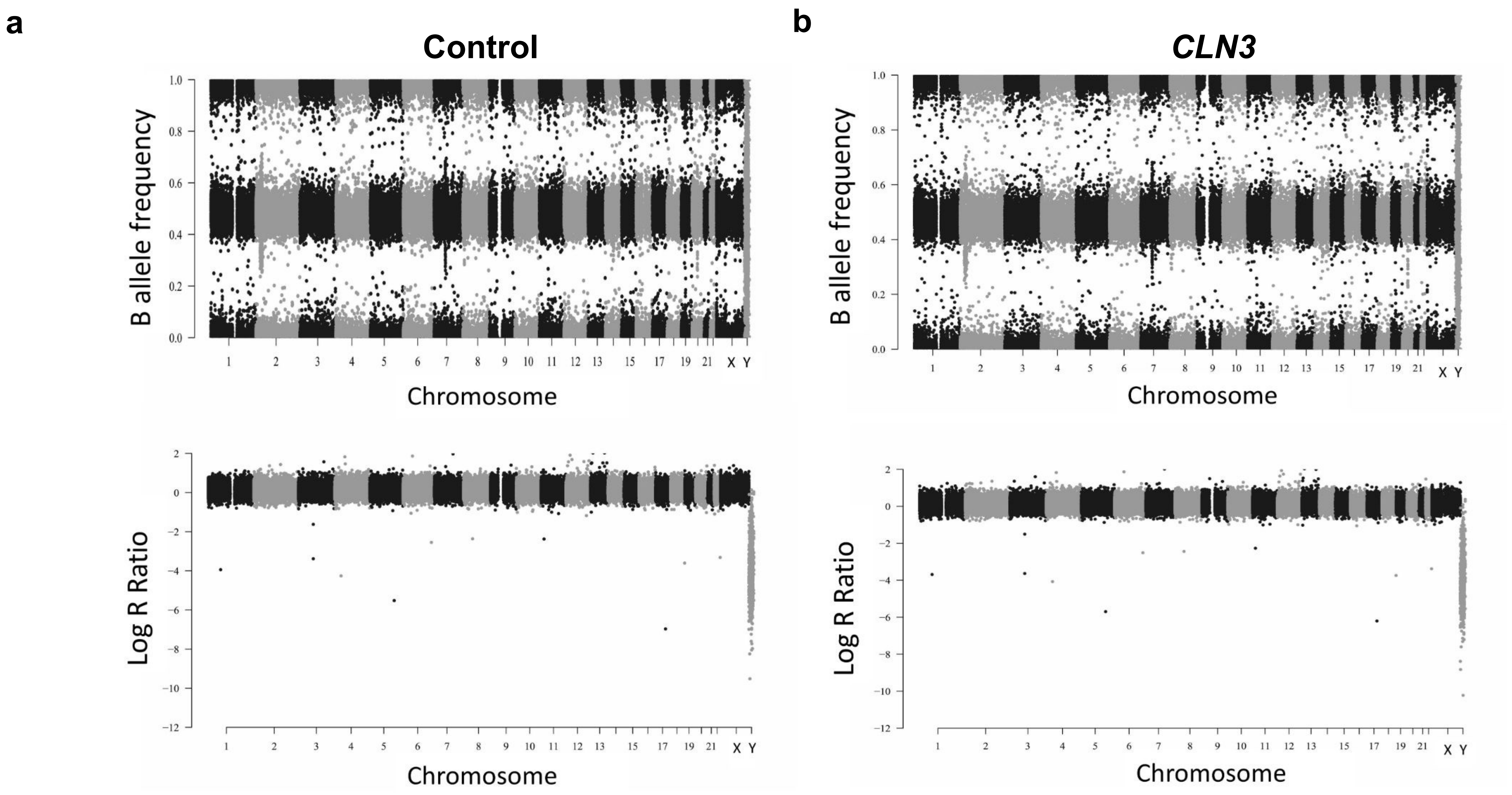

#### Supplementary Figure 2

a

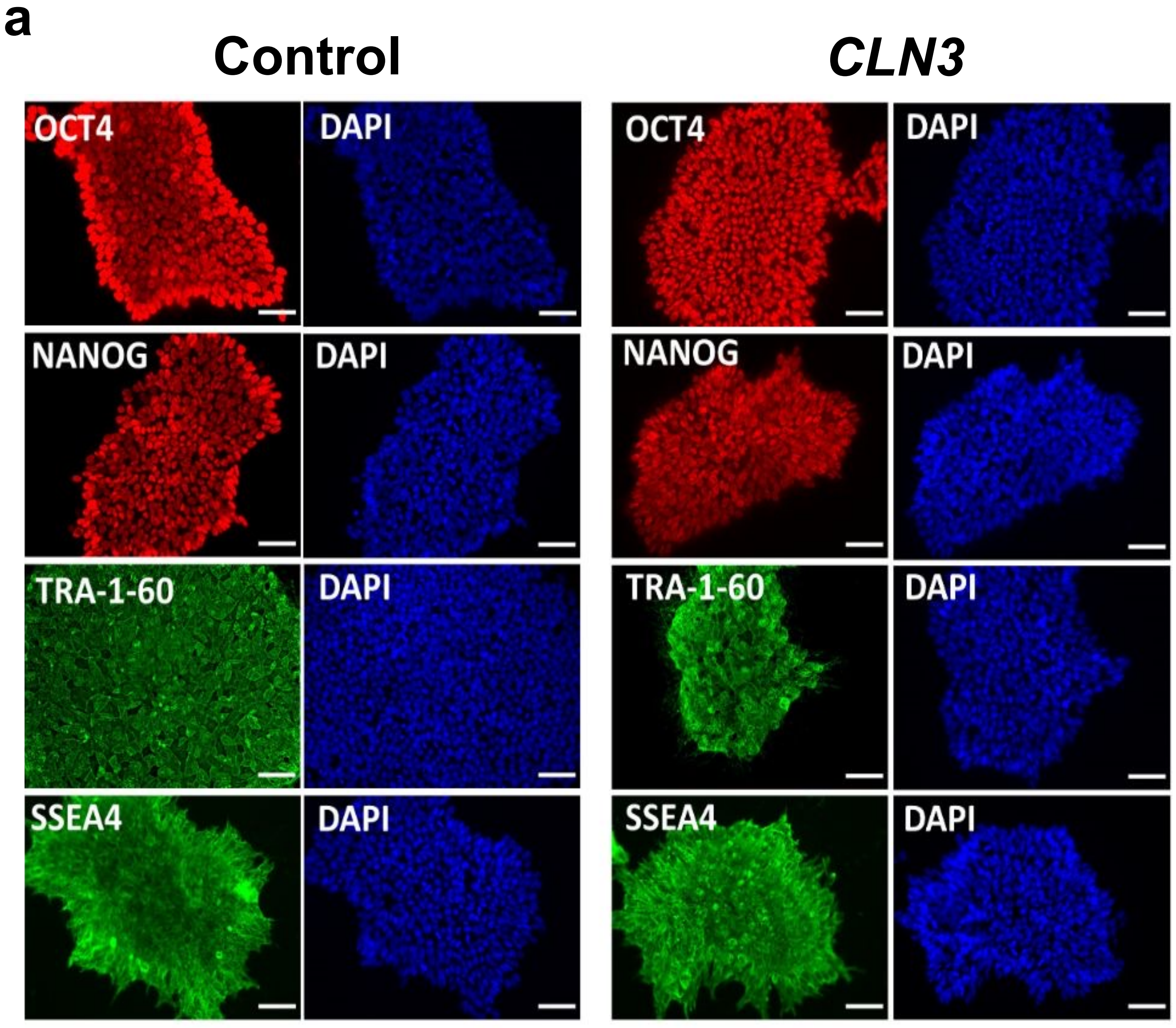


b

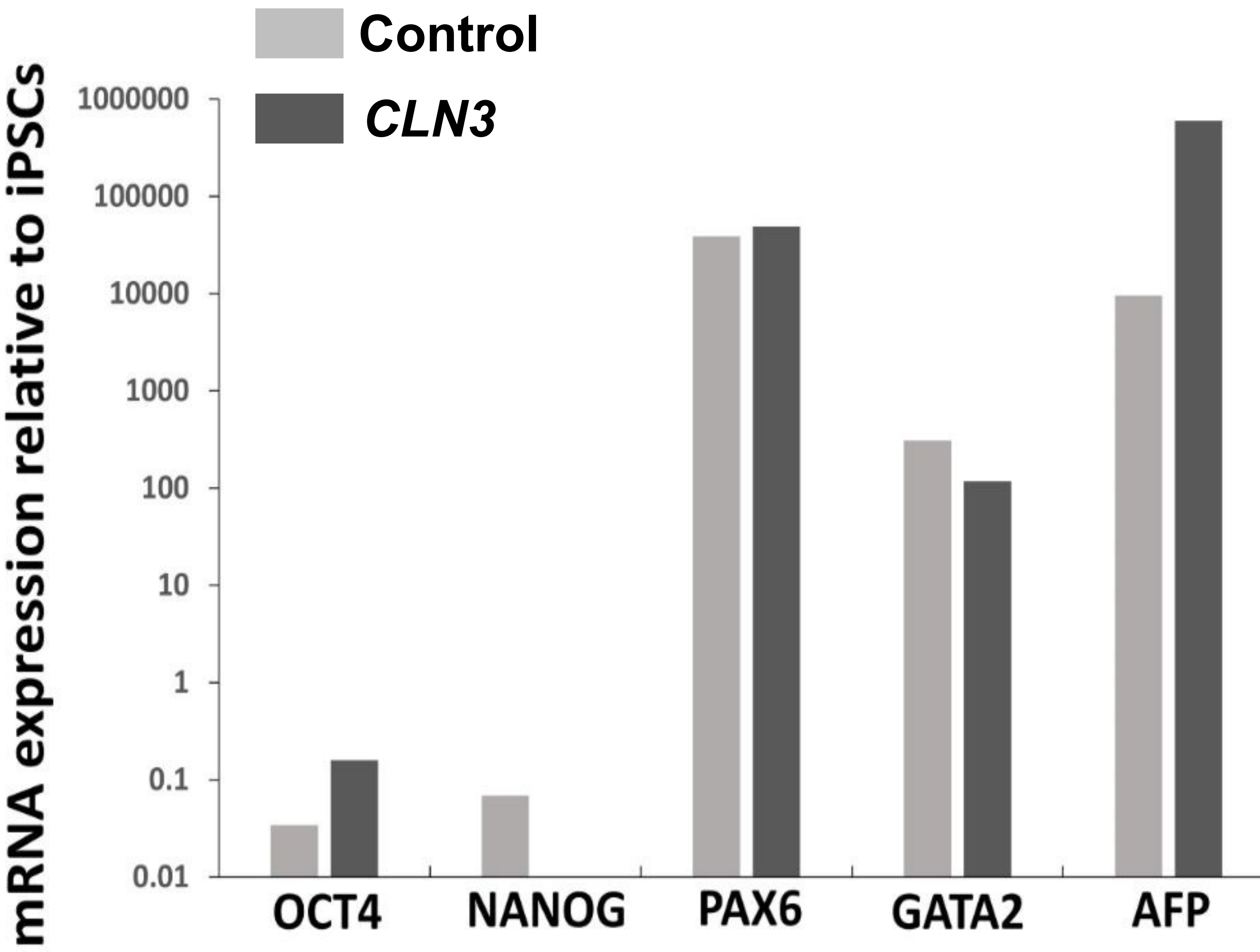

**Supplementary Figure 3**

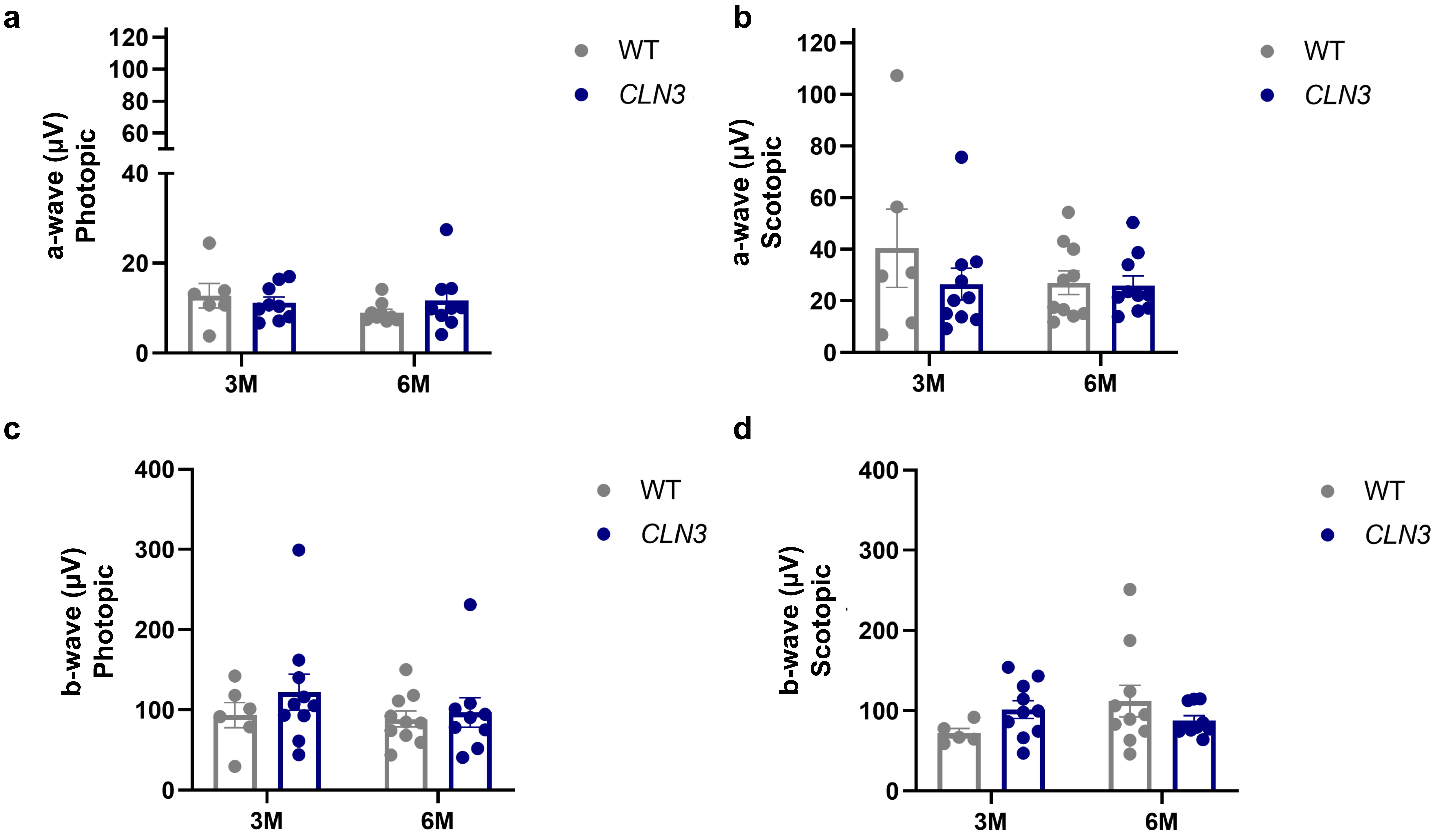

**Supplementary Figure 4**

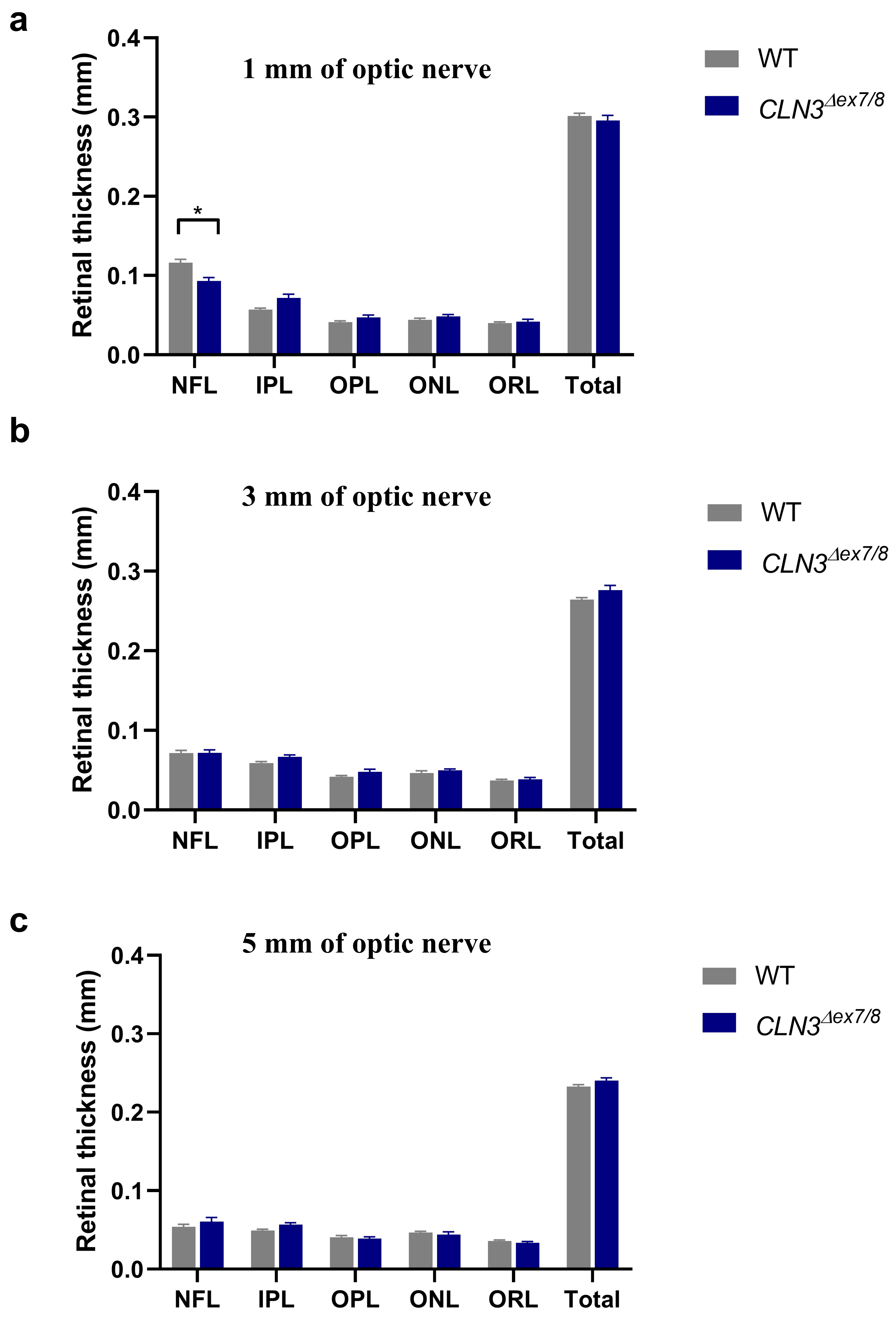
