## Supplementary material for "Genetic and cellular basis of impaired phagocytosis and photoreceptor degeneration in CLN3 disease": Table 1

**Table 1. List of CRISPR guides and primers used to confirm deletion of exons 7 and 8 in CLN3 gene**

| Oligo name | Sequence | Expected PCR band size |
| --- | --- | --- |
| sg1 | ATGAAGGGGCAGAGACATCA | NA |
| sg2 | CCTCCCTTCACAGCAAGGTA | NA |
| CLN3 F1 | GGATGAATTAGATGGAGATTGAGG | Deleted: 600-800 bp  Undeleted: 1529bp |
| CLN3 R1 | CTCATCCTACTTCTAATCACCTTG |  |
| CLN3 F2 | TCTGTCTCTACGGCTGCTGTGC | Deleted: 578bp  Undeleted: 795bp |
| CLN3 R2 | GAACACCAGGTTGAGGCACTGC |  |
