## Supplementary material for "Genetic and cellular basis of impaired phagocytosis and photoreceptor degeneration in CLN3 disease": Table 2

| **Antibody** | **Manufacturer** | **Cat. #** | **Dilution**  **Immunocytochemistry** | **Dilution (WB)** |
| --- | --- | --- | --- | --- |
| ACTN | Cell Signaling | 4970 |  | 1:500 |
| ACTN | Santa Cruz | SC-47778 |  | 1:500 |
| BEST1 | Millipore | MAB5466 |  | 1:500 |
| CRALBP | Abcam | AB15051 |  | 1:500 |
| EZR | Cell Signaling | 3145S | 1:100 | 1:1000 |
| MERTK | Abcam | AB52968 |  | 1:500 |
| RHO | Millipore | MABN15 |  | 1:500 |
| OCT4 | Cell Signaling Technology | StemLight™ Pluripotency Antibody Kit #9656 | 1:200 |  |
| NANOG | Cell Signaling Technology | StemLight™ Pluripotency Antibody Kit #9656 | 1:200 |  |
| RPE65 | Millipore | MAB5428 |  | 1:500 |
| TRA-1-60 | Cell Signaling Technology | StemLight™ Pluripotency Antibody Kit #9656 | 1:200 |  |
| SSEA4 | Cell Signaling Technology | StemLight™ Pluripotency Antibody Kit #9656 | 1:200 |  |
| ZO1 | Life Technologies | 61-7300 | 1:100 |  |
| ZO1 | Invitrogen | 33-9100 | 1:100 |  |
